# Analysis of Protein-Protein Interactions networks and cross-species transfer learning comparison for seven organisms

**DOI:** 10.1101/2023.06.05.543725

**Authors:** Yasmmin C Martins

## Abstract

**Motivation:** Protein-protein interactions (PPIs) can be used for a plenty of applications like inferring protein functions or even helping the drug discovery process. For human specie, there is a lot of validated information and functional annotations for the proteins in its interactome. In other species, the known interactome is much smaller compared with human and there are many proteins with few or no annotations by specialists. Understanding the interactome of other species helps to trace evolutionary characteristics, compare important biological processes and also build interactomes for new organisms according to other organisms more related with it instead of relying just to the human interactome.

**Results:** In this study, we evaluate the performance of PredPrIn workflow in predicting interactome for seven organisms in terms of scalability and precision showing that PredPrIn gets over than 70% of precision and it takes less than three days even on the largest datasets. We made a transfer learning analysis predicting an organism interactome from each other organism, we then showed an implication regarding to their evolutionary relation in the number of ortholog proteins shared between these organisms. We also present an analysis of functional enrichment showing the proportion of shared annotations between positive and false interactions predicted and extraction of topological features of each organism interactome such as proteins acting as hubs and bridge between modules. From each organism, one of the most frequent biological processes was selected and the proteins and pairs present in it were compared in terms of quantity in the interactome available in HINT database for that organism and the one predicted by PredPrIn. In this comparison we showed that we covered those proteins and pairs covered in HINT and also enriched these processes for almost all organisms.

**Conclusions:** In this work, we have proved the efficiency of PredPrIn workflow for protein interaction prediction for seven different organisms using scalability, performance and transfer learning analyses. We have also made cross-species interactome comparisons showing the most frequent biological processes for each organism as well as the topological features of each organism interactome showing the consistency with hypothesis about biological networks. Finally, we described the enrichment made by PredPrIn in selected biological processes showing that its prediction was important to enhance information about these organisms interactomes.

## Introduction

Protein-protein interaction (PPI) prediction has been the target of research in the last years because it provides massive information about protein networks with low cost in contrast with experimental techniques [1]. PPIs enables the understanding and investigation of important features in a specific organism such as protein function assignment and signalling pathways.

The analysis of PPIs from the network topology perspective reveals insights about the role and functions of some proteins nodes. The human PPI contains an amount of very well annotated proteins and it is used as reference to build networks for other organisms using orthologous proteins between the source (human) and target organism [2, 3]. Not all organisms have the amount of validated information such as the human and it is important to evaluate predictions for other organisms to enrich the protein network information about other species [4].

In this paper, we evaluated PredPrIn workflow for protein interaction prediction in terms of scalability and classification performance using data from HINT [4] and STRING [5] databases about seven organisms such as *Homo sapiens, Arabidopsis thaliana, Caenorhabditis elegans, Drosophila melanogaster, Escherichia coli, Mus musculus* e *Saccharomyces cerevisiae*. We compared the total execution times and precision values along six different sizes of datasets with [6], showing PredPrIn’s efficiency for all these organisms.

In order to evaluate and explore the evolutionary relations, we tested the transfer learning between the protein networks [7, 8] of these organisms and we found that most of the predictions showed precision values higher than the cutoff value of 0.7. Then, we explored the number of proteins and protein interactions shared by each pair combination of organisms and compared with the cladogram of these species showing that closer organisms in terms of evolution tends to share more orthologous proteins and pairs.

Finally, we have made functional enrichment analysis in the protein networks for all organisms. In this analysis, we investigate the Gene Ontology [9] terms sharing and the role of proteins acting as hubs and connexion between modules. We also compared the information gain with PredPrIn prediction in selected biological processes. These analysis showed that we could improve the amount of information about these organisms interactome.

## Methods

### Data acquisition

We collected high confidence binary interactions from the HINT database [4] as part of the positive portion and the other part was extracted from STRING [5] (with scores between 700 and 1000). The interactions considered false were collected from STRING with scores between 100 and 200. This data was retrieved for the following organisms: *Homo sapiens* (*H. sapiens*), *Arabidopsis thaliana* (*A. thaliana*), *Caenorhabditis elegans* (*C. elegans*), *Drosophila melanogaster* (*D. melanogaster*), *Escherichia coli* (*E. coli*), *Mus musculus* (*M. musculus*) e *Saccharomyces cerevisiae* (*S. cerevisiae*).

### Datasets construction for multiorganism prediction evaluation

We designed the datasets from the aforementioned reference databases according using six different sizes containing 5000, 10000, 25000, 50000, 100000 and 200000 pairs, with 50% of positive (high confidence score) and 50% of negative pairs (low score). We evaluated all these datasets for the seven organisms using rounds of the same size for all organisms at the same time. So, the first experiment had the size of 200 thousand for all organisms and the last had the size of 5 thousand.

The goal of these sizes was testing the precision and scalability by measuring the total execution time. We also divided the experiments in two parts, the first part we did not use the knowledge base created in the first step of PredPrIn ([10]), all the proteins information was recreated for all the sizes. In the second part, we reused the proteins information that was previously processed. The goal of using these two parts is testing the efficiency of the knowledge base construction in the first step of PredPrIn. We formed the base reused in the second part merging the information obtained in the first part for all the sizes and storing it, temporally, in another location.

### Multiorganism interactome prediction

The interactome of each organism was generated using positive predicted interactions provided by running the round with 200 thousand sizes. We used the workflow PredPrIn ([10]) for protein interaction prediction, which has three steps: (i) Protein information acquirement for all proteins involved in the pairs given for evaluation as input; (ii) Features generation using six detection methods, semantic similarity for the Gene Ontology [9] terms annotated (it was used for the three branches (Molecular function, cellular component and biological process)) for the proteins by Pekar metric [11]; analysis of domain interactions [12] corresponding to a structural type of method; co-occurrence in metabolic pathways from KEGG [13] and the last method is SPRINT [14], that is based on the primary sequence of amino acids; (iii) Classification and analysis of numerical feature vectors representing the pairs and exportation of the performance evaluation results.

These three steps generated the functional annotations that were used to do the enrichment analysis and we filtered the input pairs using the final score given by the third step to generate the protein network for each organism.

### Gene ontology analysis

For this analysis, the most frequent biological processes were found using the following method: check all different biological process annotated for all proteins involved in the pairs and then counting the amount of occurrences of each process in the pairs.

Other analysis involving functional enrichment was finding the difference between the number of pairs and proteins that PredPrIn added in some representative biological process chosen for each organism. We executed this analysis, separating two protein networks for each organism: one with the pairs shared by PredPrIn and HINT and other with only PredPrIn predicted positive pairs. Using these two networks, we filtered the pairs whose proteins were annotated with the chosen biological process for the organism and count the amount of different proteins and pairs remaining in the two networks after selection.

## Results and discussions

In this section, we present the analysis made considering seven organisms, which are: *Homo sapiens* (*H. sapiens*), *Arabidopsis thaliana* (*A. thaliana*), *Caenorhabditis elegans* (*C. elegans*), *Drosophila melanogaster* (*D. melanogaster*), *Escherichia coli* (*E. coli*), *Mus musculus* (*M. musculus*) and *Saccharomyces cerevisiae* (*S. cerevisiae*). These analysis comprised an strategy to tackle the computational validation for multiple organisms of PredPrIn workflow and describe characteristics of the predicted interactome of each organism. In this context, the following evaluation criteria were used: (i) Computational validation with performance, scalability and comparison with the related work [6]; (ii) Transfer learning between the organisms predictions and (iii) Functional enrichment with confirmation of biological hypothesis, identification of frequent biological processes and case studies of enriched process with PredPrIn in each organism.

### Performance and scalability prediction evaluation

The first part of the computational validation was evaluating the performance using the precision metric and its distribution for the organisms according to the different dataset sizes (as shown in figure 1).

**Figure 1.**
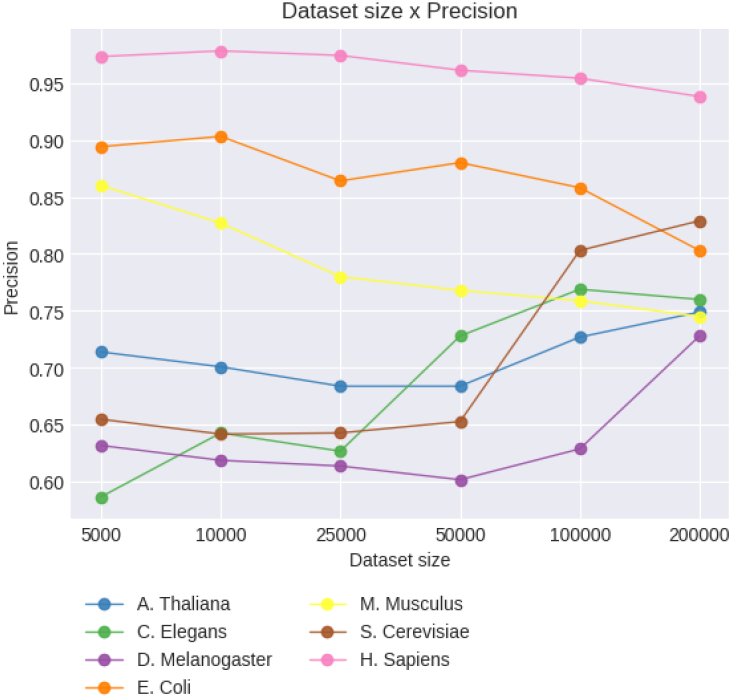
Distribution of the precision values according to the dataset sizes for each organism

In this figure, it is possible to observe that the organisms *M. musculus, E. coli* e *H. sapiens* had a good performance (above 0.75) since the smaller dataset and the precision values started to decrease after the size of 50000. *M. musculus* had a major decrease (from 0.84 to 0.77) between 10000 and 25000 and then the values stabilised. One reason for this precision declining for *E. coli* and *M. musculus* between 10000 and 25000 is that the positive examples from HINT (high confidence) finished and the positive portion was completed with STRING data with high score, which may have proteins with high or low annotation quality. Meanwhile, the other four organisms started with precision values between 0.58 and 0.73 and gradually increased the precision according to dataset size. Overall, PredPrIn got high values for precision and for some organisms the precision value was increased with higher sizes of datasets. This shows that PredPrIn can learn and improve its performance not only for human genome but also for other organisms.

In [6], a similar analysis was made using training dataset size and precision values. In the mentioned work, they obtained six datasets with distinct sizes (1000, 2500, 5000, 10000, 25000 and 50000) and the pairs were also obtained from HINT. The precision in 5000 is around 0.5 and in 50000 it increases to 0.7, while in the experiment using PredPrIn, in 5000 all precision values is around 0.60, and in 50000, four organisms had 0.7 or more and in 100000 only one (*D. melanogaster*) does not increased this value, but it had a significant raise (from 0.63 to 0.73) for 200000 size.

Other evaluation made was for scalability, measuring the total execution time of PredPrIn for distinct dataset sizes in each organism with and without the protein knowledge base generated along the experiments execution (the results can be observed in figure 2).

**Figure 2.**
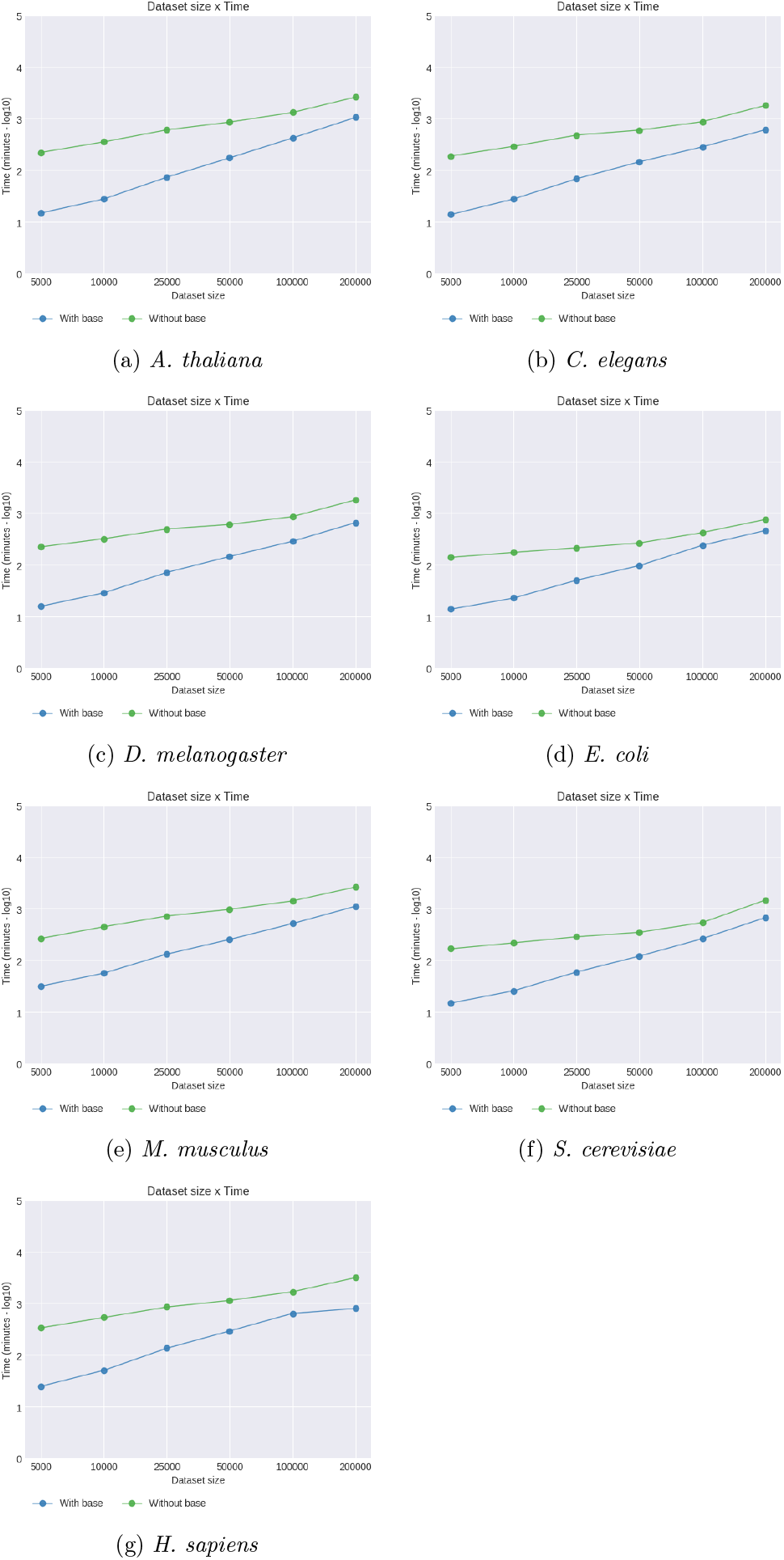
Distribution of the execution time (minutes) along the dataset sizes for each organism.

The plots contain the total execution time, including data acquirement, numerical features calculation and pairs classification. As expected, the time spent to execute PredPrIn workflow increases according to the size of the datasets. The highest execution time was found for the organism *H. Sapiens* (3000 minutes, approximately), which has the biggest number of different proteins and most of them has several functional annotations. Other pattern observed is that for the sizes 5000, 10000 an 25000, the difference of time between the experiments with and without use of the knowledge base is bigger than the difference in the bigger datasets, despite the fact of still have an advantage of time reduction. The difference reduction is explained by the fact of the features calculation step take as much time as the number of pairs. Because of this, even optimising the first step, the second step remains computational expensive.

[6] (DPPI) also did an analysis relating dataset size and time. The biggest dataset for this work (50000) spent around 1000 minutes, the most time-consuming dataset for PredPrIn (belong to *H. Sapiens* organism) with 50000 also spent a similar amount of minutes. However, the mentioned work considers only the training time and not the total execution time. DPPI, needs an extensive data pre-processing step till obtaining the data to feed the deep neural network proposed by the method. So, if we count with this step, PredPrIn is more efficient because it spent the same time even counting the data acquisition and preparation step.

Another analysis was made to investigate the execution time spliced for each step of PredPrIn workflow (in figure 3) with the goal of finding points that can be improved and optimised in the future, taking less time and being more efficient.

**Figure 3.**
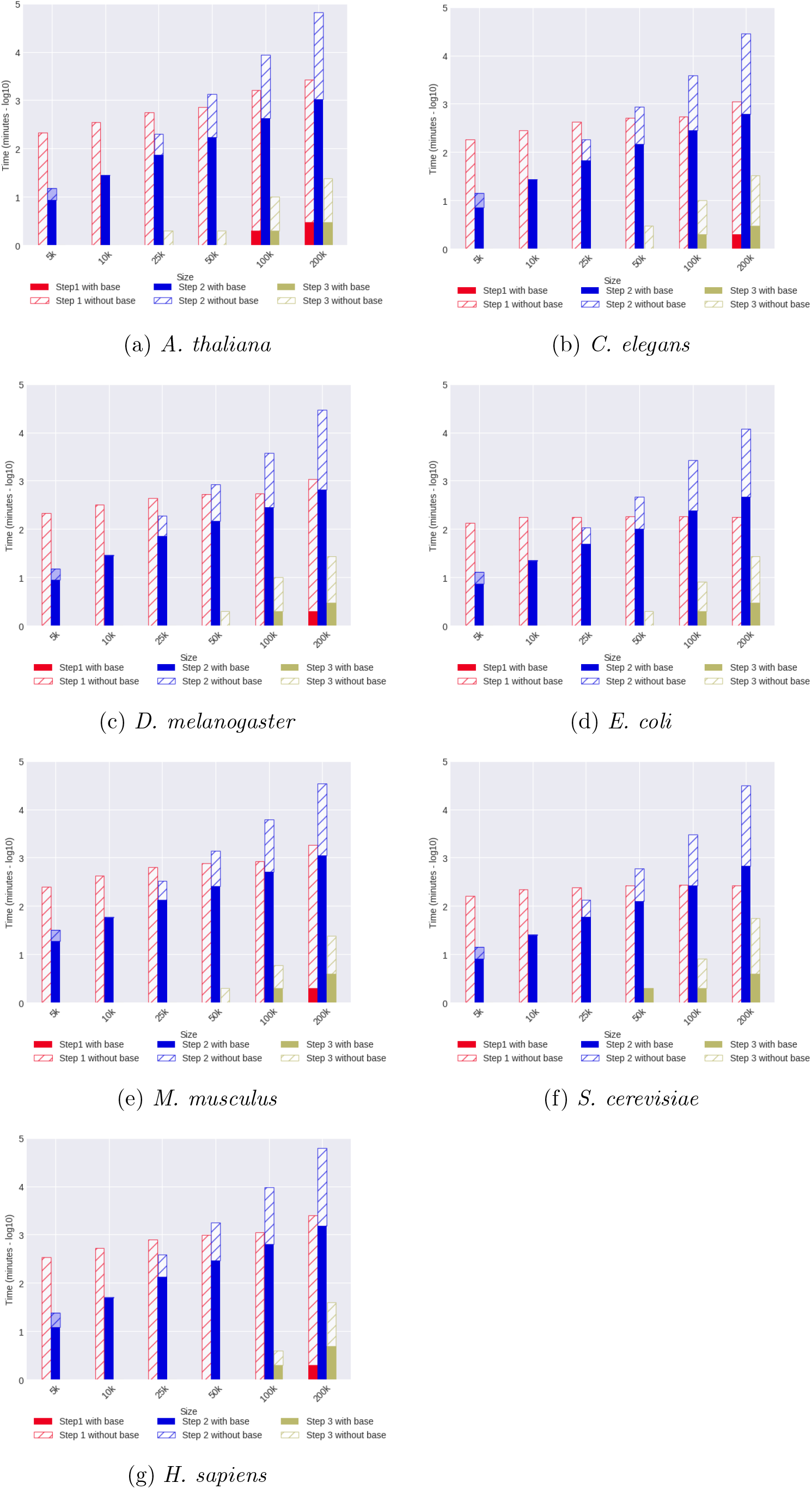
Execution time of each PredPrIn step in the diverse dataset sizes organized by organism.

We can observe that the knowledge base construction reduced almost 100% of the first step time cost for almost all datasets. Furthermore, the longest execution times belong to the organisms *H. sapiens, M. musculus* and *A. thaliana*, and the most expensive step is the pre-processing and the second is the features generation. This second step makes the probability calculation according to the different detection methods and it does not change significantly its performance with the knowledge base usage. Three of the six features calculated in this step are provided by semantic similarity, and the annotations sets for each protein are combined using the *Best-Match Average* [15] technique that spends some amount of seconds to be computed and it is proportional to the quantity of annotations for each protein. Some manners of decreasing the impact is testing the performance of other techniques. The step responsible for classifying the pairs is the most cheap, taking less than 15 minutes even in the biggest datasets.

### Transfer learning analysis

The analysis contained in this section have as goal verifying the transfer learning ability between a model trained in one organism to test data of other organisms. Transfer learning can be defined as the usage of data knowing its labels to predict labels using unlabeled data. This analysis was made by some works under different aspects of protein interactions [7, 8].

The organisms used to execute the analysis are those representative from different hierarchies of the species evolutionary line and, in this section, we excluded the *H. Sapiens* organism since there already is another representative for mammals (*M. musculus*). From the experiments made in the previous section, we selected the trained models of datasets with 200000 pairs and the datasets of the other 5 organisms were used as test set to evaluate the model. The prediction results are described in table 1.

**Table 1.** Table with precision values of prediction using a model of one organism testing it with dataset of other 5 organisms.

| Modelo/Alvo | *A. thaliana* | *C. elegans* | *D. melanogaster* | *E. coli* | *M. musculus* | *S. cerevisiae* |
| --- | --- | --- | --- | --- | --- | --- |
| *A. thaliana* | - | 0.809 | 0.699 | 0.645 | 0.746 | 0.799 |
| *C. elegans* | 0.729 | - | 0.682 | 0.599 | 0.741 | 0.771 |
| *D. melanogaster* | 0.783 | 0.829 | - | 0.645 | 0.782 | 0.830 |
| *E. coli* | 0.834 | 0.858 | 0.790 | - | 0.815 | 0.883 |
| *M. musculus* | 0.782 | 0.793 | 0.716 | 0.676 | - | 0.829 |
| *S. cerevisiae* | 0.785 | 0.782 | 0.716 | 0.670 | 0.756 | - |

We can observe in this table that a great portion of the values was high (above 0.75) and the worst values (lower than 0.7) were found for all models tested in the *E. Coli* dataset and also testing the model of *A. Thaliana* and *C. Elegans* in *D. melanogaster* dataset. These results indicate that PredPrIn can learn features of one organism model and use this information for other organisms data. This is important to infer protein interactions in genomes [3, 16] whose interactomes are still not available in reference databases.

We also made an orthology study showing the relation between these organisms by discovering their shared proteins and interactions. For this study, we used the mapping files provided by Uniprot database containing the identifier and the other auxiliary label which is formed by the name in HGNC (HUGO Gene Nomenclature Committee) format concatenated to the organism identifier, following this scheme: _HUMAN for *H. sapiens*, _ARATH for *A. thaliana*, _CAEEL for *C. elegans*, _MOUSE for Mus Musculus, _DROME for *D. melanogaster*, _ECOLI for *E. coli* and _YEAST for *S. cerevisiae*. Using this auxiliary identifier is possible doing intersections between sets of pairs and proteins of each organism. Table 2 shows the number of proteins and pairs that were conserved between the organisms interactomes. The pairs used to calculate the intersections are those that were predicted as positive by PredPrIn in the experiments with 200000 pairs.

**Table 2.** Table with the number of proteins and interactions shared between the organisms (# proteins/# pairs).

| Org./Org. | *A. thaliana* | *C. elegans* | *D. melanogaster* | *E. coli* | *M. musculus* | *S. cerevisiae* |
| --- | --- | --- | --- | --- | --- | --- |
| *A. thaliana* | - | (556/150) | (468/124) | (168/47) | (1166/205) | (895/423) |
| *C. elegans* | (556/150) | - | (825/1068) | (130/385) | (1202/980) | (729/944) |
| *D. melanogaster* | (468/124) | (825/1068) | - | (138/267) | (1306/907) | (611/876) |
| *E. coli* | (168/47) | (130/385) | (138/267) | - | (250/311) | (210/298) |
| *M. musculus* | (1166/205) | (1202/980) | (1306/907) | (250/311) | - | (1226/1315) |
| *S. cerevisiae* | (895/423) | (729/944) | (611/876) | (210/298) | (1226/1315) | - |

The ortholog proteins are those that keep their function along the evolutionary line and despite of having the same function in distinct organisms, their primary sequences of amino acids can suffer modifications [17]. The evolutionary relationship between the species involved in this analysis can be observed in figure 4. We used the taxonomy tool from NCBI (National Center for Biotechnology Information)^[1]^ to retrieve the cladogram information about the species and we exported this information as phylip file format. Using this file, a image showing the cladogram was rendered with the visualisation tool ETE Toolkit ^[2]^.

**Figure 4.**
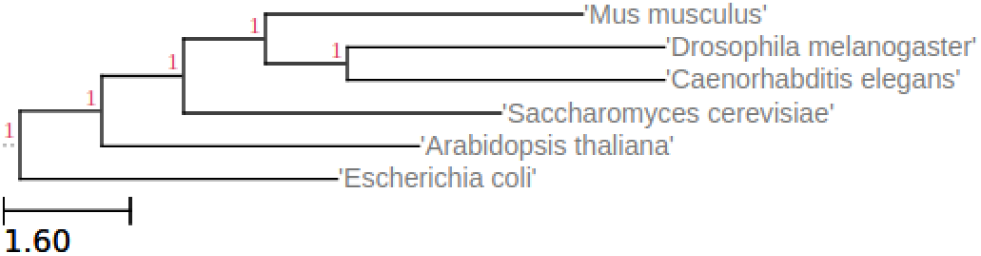
Organisms evolutive relationship.

We can notice in this table that *A. thaliana* is the one that have less proteins and pairs in common with other organisms, this makes sense with its relation among the other organisms showed in the cladogram since it is a representative of *Plantae* kingdom and this implies in several distinct biological process between this plant and the rest of the organisms [18]. The same behaviour was found for the bacteria *E. coli*, which is a procaryont organism whose genetic material is not contained in a nucleus^[3]^, sharing few proteins and interactions with the others. *M. musculus, C. elegans* and *D. melanogaster* are part of *Animalia* kingdom. and have several interactions and proteins in common. *S. cerevisiae* organism is a representative of *Fungi* kingdom and is nearest of *Animalia* kingdom participants and share more proteins and pairs with these ones than with bacteria and plant representatives. These observations confirm that the interactome provided by PredPrIn prediction represents specialities and evolutive relations amidst organisms.

### Functional enrichment and topological features of the interactomes

In this section, we present results of a functional enrichment analysis having as goals (i) confirming the hypotheses of the functional annotations sharing, (ii) finding most frequent biological processes in each organism, and (iii) choose a representative process for each organism to give details and compare proteins and pairs in HINT for these process and those proteins that were found in the interactome provided by PredPrIn.

For analysis (i), we investigated terms sharing of biological process branch of gene ontology and a more restricted case which is the terms sharing for the three gene ontology branches at the same time by the proteins involved in pairs.

In figure 5, the terms sharing plots keep the expected behaviour, which means that the pairs predicted as positive keep sharing more terms than the pairs predicted as false. The difference between the positive and negative part was significant in all datasets (the number of positive pairs sharing was twice or more in relation to the false pairs), except for *C. elegans* interactome. In this interactome, 27% of the positive pairs shared terms of biological process and 24% for the false pairs, and 10% of the positive pairs shared the terms for the three branches against 8% for the false pairs. Most part of occurrences were found for *M. musculus, S. cerevisiae* and *H. sapiens*. For these three organisms, the terms sharing of biological process branch for positive portion of the pairs were 34%, 43% and 47% and for the false portion were 19%, 10% and 12%. The terms sharing for the three branches in the positive pairs were 14%, 17% and 23%, for the false pairs were 6%, 3% and 3%, respectively.

**Figure 5.**
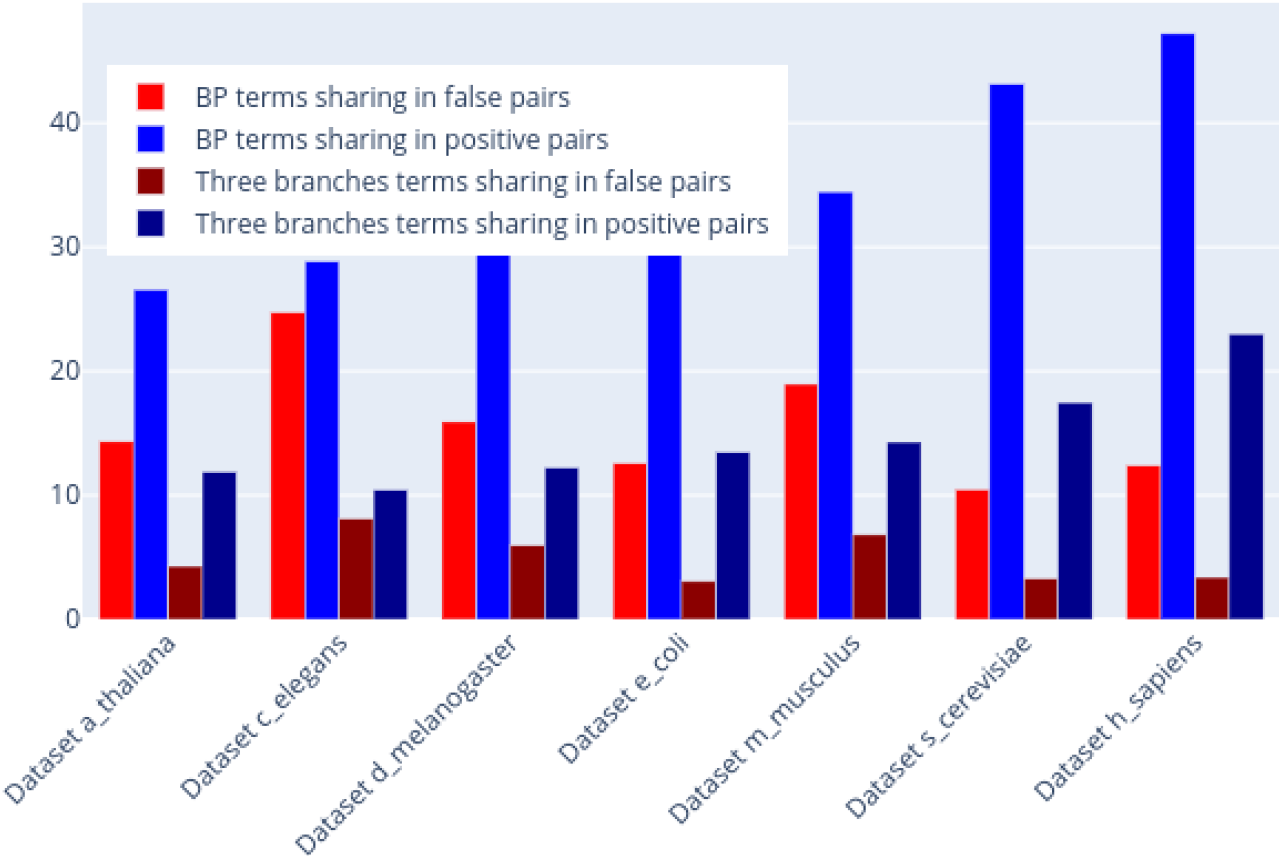
Biological process and all GO branch terms sharing for the positive and negatively predicted ppis among the organisms.

Such analysis confirmed that even in other organisms the hypotheses about as higher the functional annotations sharing is, higher is the probability of occurring interactions was confirmed.

For analysis (ii), we identified the most frequent biological processes among proteins annotations belonging to the pairs of the organisms interactomes. The goal of this analysis is showing the main processes of each organism.

Some of the most frequent biological processes found are general and common to several organisms like the translation process and *mRNA splicing, via spliceosome*, which refers to exons (coding regions) union of one or more primary transcripts of mRNA and the exclusion of intron sequences by the spliceosome mechanism and creating a mRNA consisting of united exons^[4]^.

Some particular processes were found in specific organisms, like the process *response to cadmium ion* for *A. thaliana*, which means the response to cadmium ion and it is related to the regulatory pathways of stress caused by cadmium in plants (which is the kingdom that this organism belongs to) [19]. For the bacteria *E. coli*, one of the most frequent process was the *response to antibiotic* and the antibiotics are drugs that have mechanisms to inhibit the bacterial action [20]. For *C. elegans*, we found the process of *determination of adult lifespan* and this organism is a model system to study the genetic basis of aging [21]. For the organisms *M. musculus* and *H. sapiens*, we found several common processes like positive and negative regulation of transcription by RNA polymerase II, apoptotic process regulation and protein phosphorylation.

The next and last analysis (iii) investigated the amount of proteins and pairs that were added in the biological processes of the organisms with the interactome provided by PredPrIn. We have chosen a representative process for each organism to do such analysis and we count the portion of proteins and pairs that HINT and PredPrIn have in the specific process and the portion of proteins and pairs that were found only in PredPrIn interactome.

Figure 6 shows the occurrences of proteins and pairs organised by the representative process of each organism. We chose the *mRNA splicing, via spliceosome* process for *A. thaliana, C. elegans, D. melanogaster* and *S. cerevisiae*; *Response to antibiotic* for *E. coli* ; and *Positive regulation of transcription* for *M. musculus* and *H. sapiens*. We can observe that the amount of pairs added for the processes in PredPrIn interactome was high for almost all organisms except for *H. Sapiens*.

**Figure 6.**
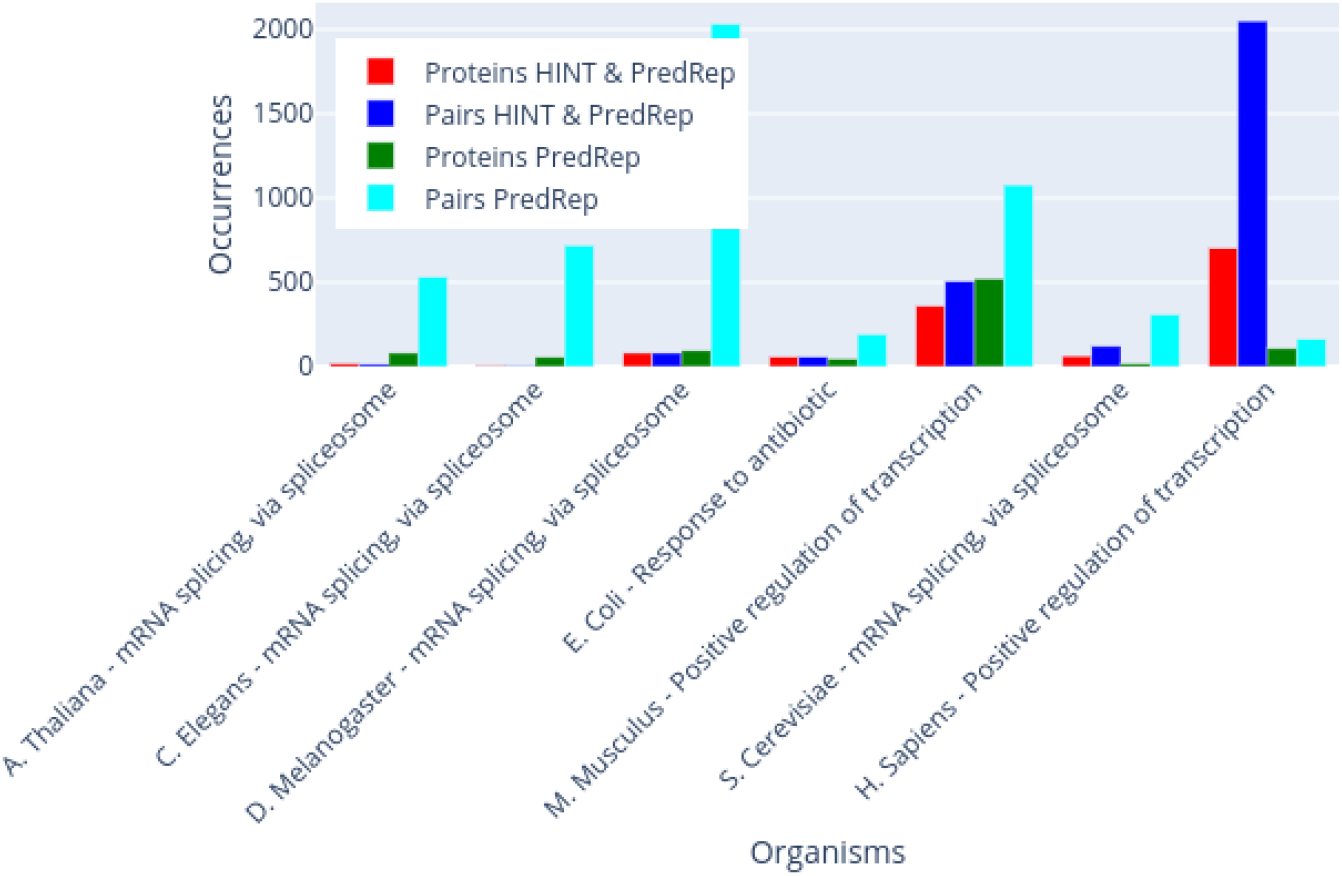
PredPrIn contribution for the representativeness increase of frequent biological processes of each organism.

*M. musculus* was the only organism whose process had a higher amount of proteins and pairs (both had numbers above 400) shared by HINT and PredPrIn. In the case of *H. sapiens*, a great part of the interactions in its process were alredy described in HINT and they were reinforced in PredPrIn, this the reason why there were so few addictions of proteins and pairs. This analysis showed the PredPrIn’s contribution in complementing the biological processes in relation to HINT interactome.

## Conclusion

In this paper, we presented an analysis of PredPrIn workflow to evaluate the scalability and performance as well as functional enrichment and transfer learning using the predicted interactomes of the organisms. The results showed that it is efficient for all the seven organisms tested and it spends less than three days to do all the steps even for datasets with 200 thousand pairs. We showed the most frequent biological process and highlighted those specific for the organisms as well as commented the process applied to proteins with important topological roles as hubs and high bettweenness. We also made a test of transfer learning predicting interactomes of one organism using a model trained in others and we show the relation between the scores obtained and the evolutionary relationship between the organisms. Finally, we selected one biological process for each organism to study the amount of proteins and pairs that PredPrIn prediction have added to the HINT interactome publicly available, showing that was a gain in almost all organisms except human because this one is very well annotated.

## Author’s contributions

Text for this section …

## Acknowledgements

Text for this section …

## Footnotes

[1] https://www.ncbi.nlm.nih.gov/Taxonomy/CommonTree/wwwcmt.cgi

[2] http://etetoolkit.org/treeview/

[3] https://www.nature.com/subjects/prokaryote

[4] http://amigo.geneontology.org/amigo/term/GO:0000398

